# CRISPR-RT: A web service for designing CRISPR-C2c2 crRNA with improved target specificity

**DOI:** 10.1101/099895

**Authors:** Houxiang Zhu, Emily Richmond, Chun Liang

## Abstract

CRISPR-Cas systems have been successfully applied in genome editing. Recently, the CRISPR-C2c2 system has been reported as a tool for RNA editing. Here we describe CRISPR-RT (CRISPR **R**NA-**T**argeting), the first web service to help biologists design the crRNA with improved target specificity for the CRISPR-C2c2 system. CRISPR-RT allows users to set up a wide range of parameters, making it highly flexible for current and future research in CRISPR-based RNA editing. CRISPR-RT covers major model organisms and can be easily extended to cover other species. CRISPR-RT will empower researchers in RNA editing. It is available at http://bioinfolab.miamioh.edu/CRISPR-RT.

## Introduction

The CRISPR (<u>c</u>lustered <u>r</u>egularly <u>i</u>nterspaced <u>s</u>hort <u>p</u>alindromic <u>r</u>epeats)-Cas (<u>C</u>RISPR-<u>as</u>sociated) system is an adaptive immune system of prokaryotes, which is used to resist foreign genetic elements in bacteria and archaea (Marraffini and Sontheimer, 2010; Barrangou and Marraffini, 2014). During the past several years, CRISPR-Cas systems have been successfully applied in genome editing (Hwang *et al.*, 2013; Jiang *et al.*, 2013; Cho *et al.*, 2013; Cong *et al.*, 2013; Bikard *et al.*, 2014; Wan *et al.*, 2015), but no systems have been reported for RNA editing. Therefore, new CRISPR-Cas systems that regulate RNA activities are necessary for studying the roles of RNA molecules. Recently, the CRISPR-C2c2 system has been demonstrated as a tool for RNA targeting (Abudayyeh *et al.*, 2016).

CRISPR-C2c2 was discovered in 21 bacterial genomes and belongs to the type VI of Class 2 CRISPR systems (Shmakov *et al.*, 2015). Researchers have characterized the CRISPR-C2c2 system from the bacteria *Leptotrichia shahii* (Abudayyeh *et al.*, 2016). The *L. shahii* C2c2 locus is simply organized, including *C2c2, Cas1, Cas2* and a CRISPR array (Figure 1). Cas1 and Cas2 play an important role in spacer acquisition (Nuñez *et al.*, 2015; Heler *et al.*, 2015; Nuñez *et al.*, 2014; Díez-Villaseñor *et al.*, 2013; Yosef *et al.*, 2012; Datsenko *et al.*, 2012). C2c2, which contains two higher eukaryotes and prokaryotes nucleotide-binding (HEPN) domains, mainly functions as a sole effector protein mediating single-strand RNA cleavage (Abudayyeh *et al.*, 2016). The CRISPR array consists of many short palindromic repeat elements and spacer sequences. Similar to other Class 2 systems, the CRISPR array of CRISPR-C2c2 is first transcribed into pre-crRNA (Shmakov *et al.*, 2015). Differently, the pre-crRNA is processed by C2c2 into mature crRNAs without attaching to trans-activating crRNAs (East-Seletsky *et al.*, 2016). The mature crRNA binds to C2c2 and guides it to target a specific single-strand RNA. Then, in the target site, the HEPN domains of C2c2 mediate cleavage of the single-strand RNA (Abudayyeh *et al.*, 2016). The crRNA stem-loop structure is also required for cleavage. Thus, the palindromic repeat length of crRNA needs to be longer than 24 nt to maintain the stem loop (Abudayyeh *et al.*, 2016). In addition, C2c2 combined with a 22-28 length of the target complementarity region of crRNA would effectively mediate cleavage (Abudayyeh *et al.*, 2016). The seed region is located in the center of the crRNA-target duplex, where is more sensitive to mismatches than the non-seed region (Abudayyeh *et al.*, 2016). Single mismatch can be fully tolerated by C2c2, but if double mismatches are located in the seed region, C2c2 is unable to cleave the single-strand RNA (Abudayyeh *et al.*, 2016). C2c2 can even tolerate 3 consecutive mismatches in the non-seed region (Abudayyeh *et al.*, 2016). Currently, there is no research about gaps (insertions or deletions) tolerated by C2c2, which may be explored in the near future. The CRISPR-C2c2 system prefers H (A, U, or C) for the 3’ protospacer flanking site (PFS) sequence of one single base length to mediate single-strand RNA cleavage (Abudayyeh *et al.*, 2016). CRISPR-C2c2 has been reprogrammed to knockdown the specific mRNA in vivo, although C2c2 may also cleave collateral RNAs (Abudayyeh *et al.*, 2016). Researchers have applied the CRISPR-C2c2 system to sensitively detect transcripts in complex mixtures (East-Seletsky *et al.*, 2016). C2c2 can be easily converted into an inactive RNA-binding protein (dC2c2) by mutating any of the two HEPN domains (Abudayyeh *et al.*, 2016). Compared to the wild type C2c2, dC2c2 has more potential applications as an RNA-binding protein (Abudayyeh *et al.*, 2016). The CRISPR-C2c2 system is opening a new chapter for the programmable CRISPR tools of RNA editing. However, until now there is no available software for designing crRNAs of the CRISPR-C2c2 system.

**Figure 1.**
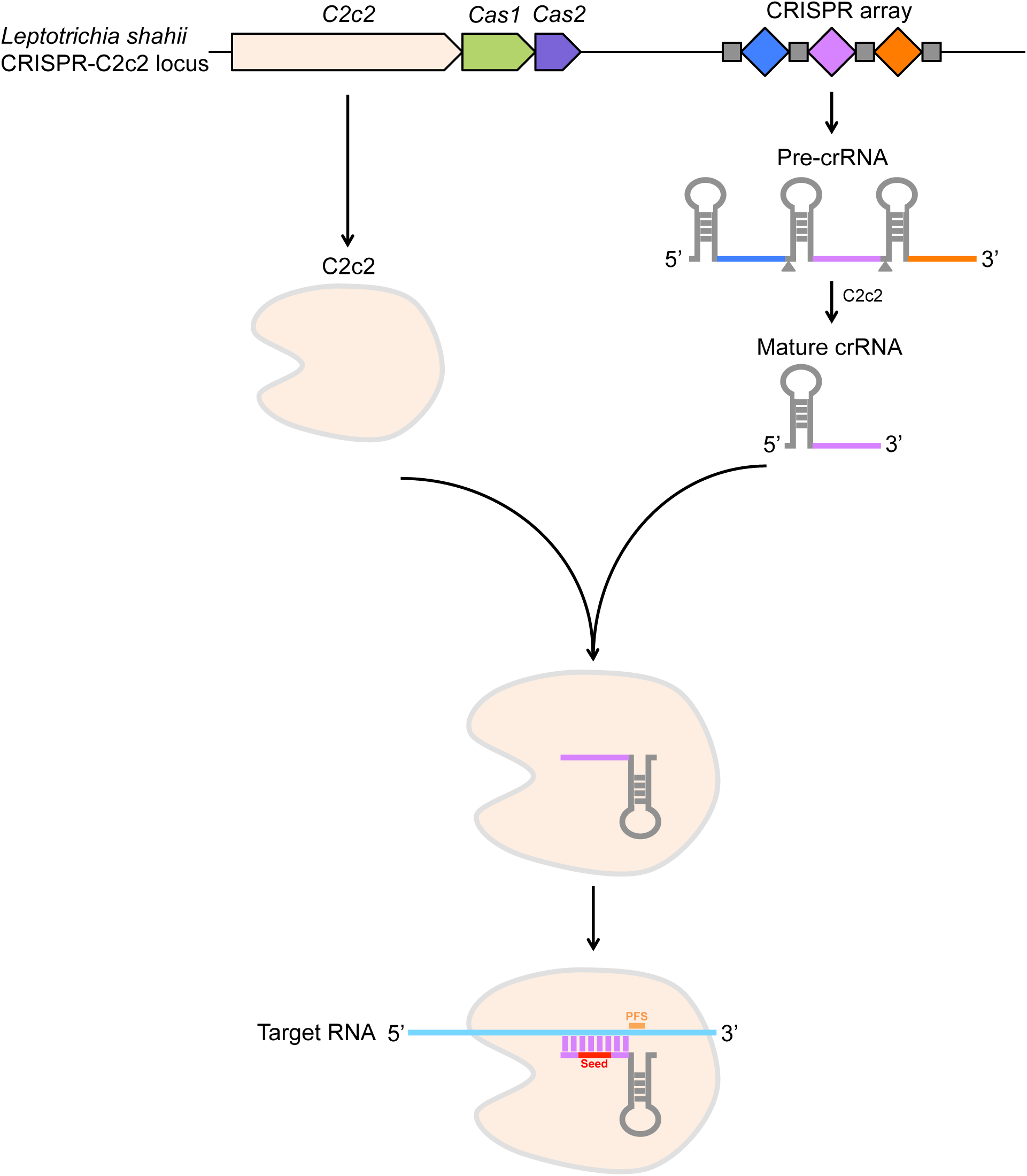
The configuration and architecture of the CRISPR-C2c2 system. The CRISPR array of CRISPR-C2c2 is first transcribed into pre-crRNA. Then, the pre-crRNA is processed by C2c2 into mature crRNAs without attaching to trans-activating crRNAs. The mature crRNA binds to C2c2 and guides C2c2 to target a specific single-strand RNA at a proper target site.

We have developed CRISPR-RT (CRISPR **R**NA-**T**argeting), a web service to help biologists design the crRNA for the CRISPR-C2c2 system. To maximize the flexibility for current and future research in CRISPR-based RNA editing, CRISPR-RT allows users to set up a wide range of parameters, such as length of the target complementarity region of crRNA, length of the seed region, the PFS, and the number of mismatches or gaps tolerated by off targets. After setting up the required parameters, CRISPR-RT will find target candidates from the input RNA sequence and employ rigorous alignment algorithms to search on- and off-target sites for each target candidate within the reference transcriptome (Langmead and Salzberg, 2012). The results are displayed in highly interactive graphical interfaces. Users can rank target candidates by the total number of target sites in the reference transcriptome, which help them choose the target candidate based on the minimum effect of off targets. In addition, users are able to validate the on- and off-target sites in the background of annotated genome and transcript features by data visualization through JBrowse (Skinner *et al.*, 2009).

## Results

### Graphic Input Interface

Figure 2 shows the home page of CRISPR-RT. If users click “C2c2” on the left side menu, the setting page of the CRISPR-C2c2 system will be displayed (Figure 3). First, users input an RNA sequence that they want to target in FASTA format into the text area. Users can also convert a cDNA sequence to the corresponding RNA sequence by clicking the “cDNA?” link. After converting, they then copy the converted RNA sequence back into the text area as the input sequence. If users want to use an example sequence, they can click the “Example Sequence” button. Second, users select a reference transcriptome. If the transcriptome that users want to choose does not show up in the drop-down menu and this transcriptome is available in ENSEMBL genome annotation (www.ensembl.org), they can click the “others?” link to submit a requirement so that the transcriptome will be added to the web server manually soon. Third, in terms of our current understanding of the CRISPR-C2c2 system architecture (Figure 1), users can set up the PFS sequence and crRNA requirements for the CRISPR-C2c2 system properly. The on- and off-target PFS sequence can be set respectively, which accepts any single letter or combination of the international union of pure and applied chemistry (IUPAC) nucleotide codes except “T” and “Z”. Users then set up the length of the target complementarity region of crRNA and the length of the seed region. The seed region is located in the center of the crRNA-target duplex, and its length should not be greater than the length of the target complementarity region of crRNA. Fourth, users can choose an off-target setting (“Basic settings” or “Specific settings”). For “Basic settings”, the number of mismatches or gaps tolerated by off targets as well as by the seed region can be set respectively. The number of consecutive mismatches or gaps in the seed or non-seed region tolerated by off targets can also be configured. For “Specific settings”, users can set up more detailed parameters in the seed and non-seed region separately, such as the number of mismatches in the seed or non-seed region tolerated by off targets. Users can also set up the search sensitivity of Bowtie2 (Langmead and Salzberg, 2012), which is related to the alignment options setting of Bowtie2, to search target sites in the reference transcriptome. Higher sensitivity setting causes alignments to be more sensitive, but it usually results in a slower search time. CRISPR-RT maximizes flexible and adjustable settings to help biologists conduct current and future experiments with more in-depth understanding of the CRISPR-C2c2 system. After setting up all the parameters, users click the “Find targets!” button which will run programs in the background to get the results. If users want to retrieve a recent job, they can click “Retrieve Jobs” on the left side menu to enter the “Job ID”, which is generated in the result page, for retrieving results in later times. The results will be kept in our server for only one week and will be deleted automatically afterwards.

**Figure 2.**
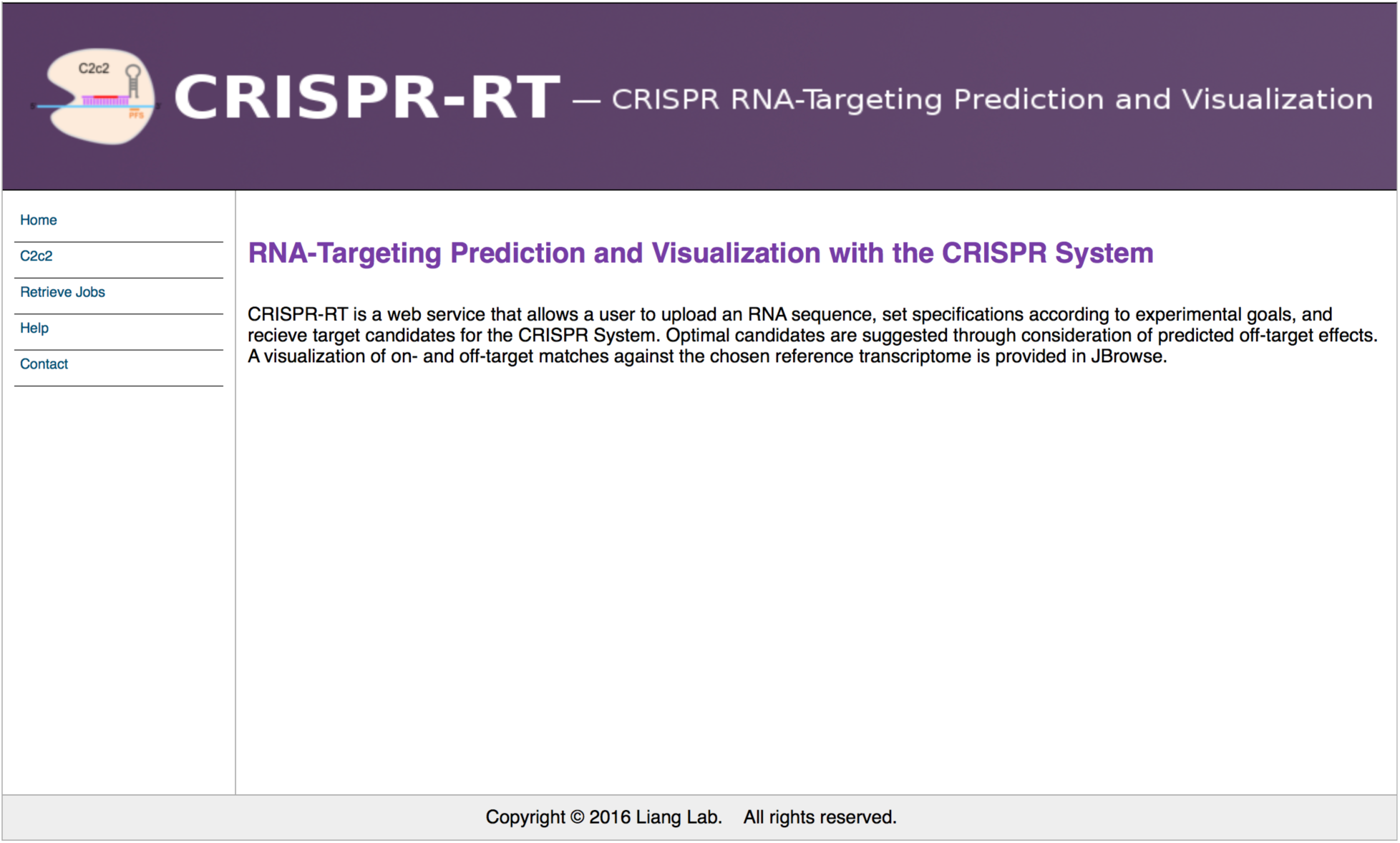
The home page of CRISPR-RT web service.

**Figure 3.**
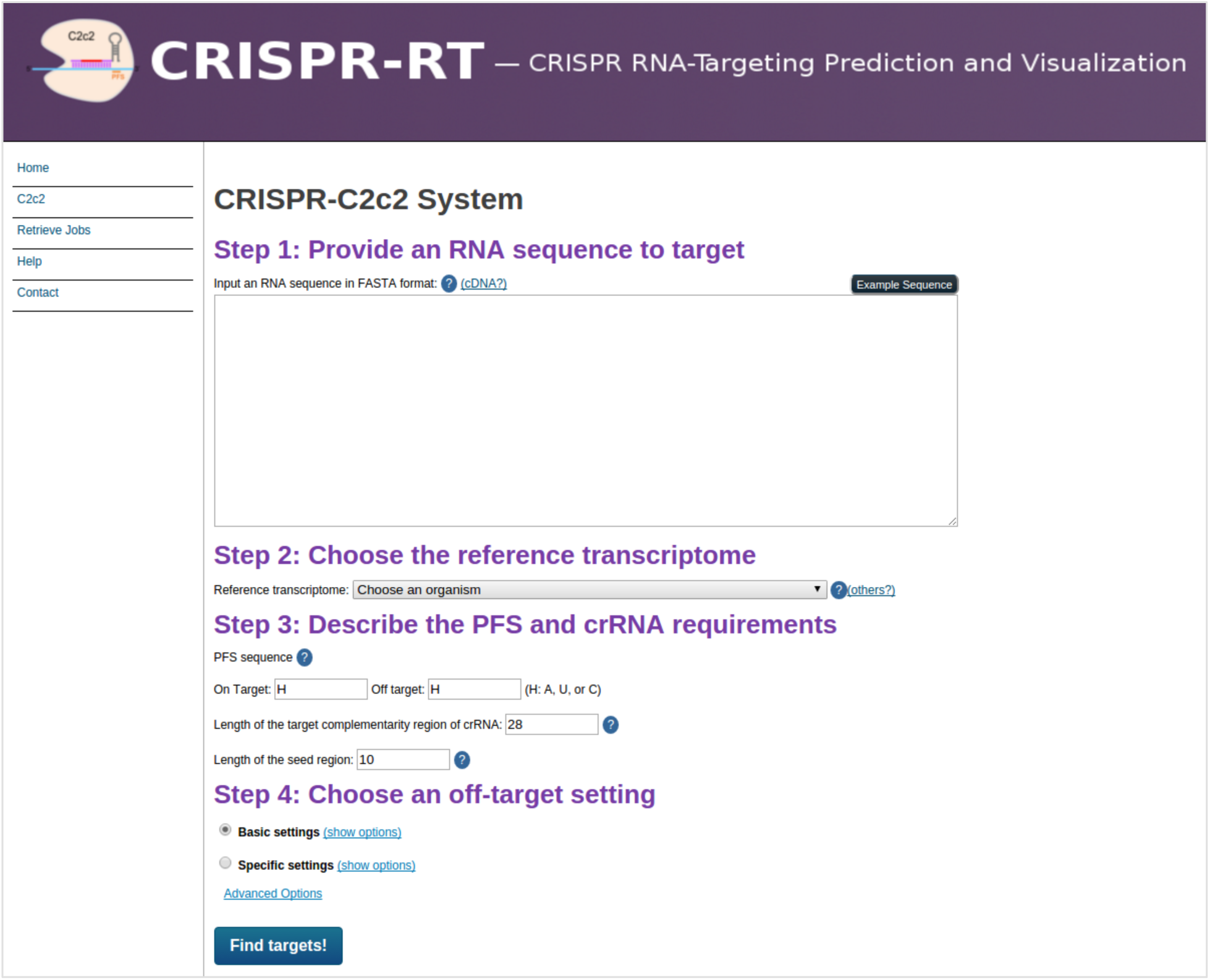
The parameter setting page of CRISPR-C2c2. First, users input an RNA sequence in FASTA format into the text area. Second, users select a reference transcriptome. Third, users set up the PFS and crRNA requirements for the CRISPR-C2c2 system. Fourth, users choose an off-target setting (either “Basic settings” or “Specific settings”).

### Graphic Output Interface

Figure 4A shows all the target candidates and relevant information for the user input RNA sequence. Users can view the input sequence by clicking the “Input Sequence Viewer” button. In the sequence viewer, users can search and highlight any subsequence such as the target candidate sequence (Figure 4B). The “Job ID” can be used to retrieve users’ recent jobs from CRISPR-RT and all relevant data will be stored for a week in our server. Users can also download the target candidates file by clicking the “Download” link. In the table, the protospacer and PFS of each target candidate are labeled by different colors. The corresponding crRNA of each target candidate can be accessed by clicking “crRNA”; a graph will appear to help users design their own crRNAs (Figure 5). The table also displays the start position, end position, and GC content of each target candidate sequence. The last column of the table shows the number of target sites in the transcriptome for each target candidate. By clicking the “#Targets” header users are able to rank all the target candidates based on the number of target sites. Target candidates with fewer number of target sites have higher target specificity in the transcriptome. If the number of target sites is 1, the corresponding table cell will be highlighted with green background color to indicate that the target candidate is highly specific in the whole transcriptome. By clicking the number of target sites users can view the detailed information of target sites for each target candidate (Figure 4C). Because CRISPR-RT has converted the transcriptome mapping result to a genome mapping result, the information of target sites are displayed in genomic context with genomic coordinates and gene annotations, including mapped gene, chromosome, strand, start/end position, and transcript isoforms. Users can click the “transcript ID” link or “gene ID” link of each target site to view the detailed description of transcript or gene where the target site located. The number of mismatches or gaps in each target site is also shown in the table. To visualize and manually validate the target sites, users can click the “JBrowse” link to visualize each target site in the background of genome and transcript features annotated by ENSEMBL (www.ensembl.org) (Figure 4D).

**Figure 4.**
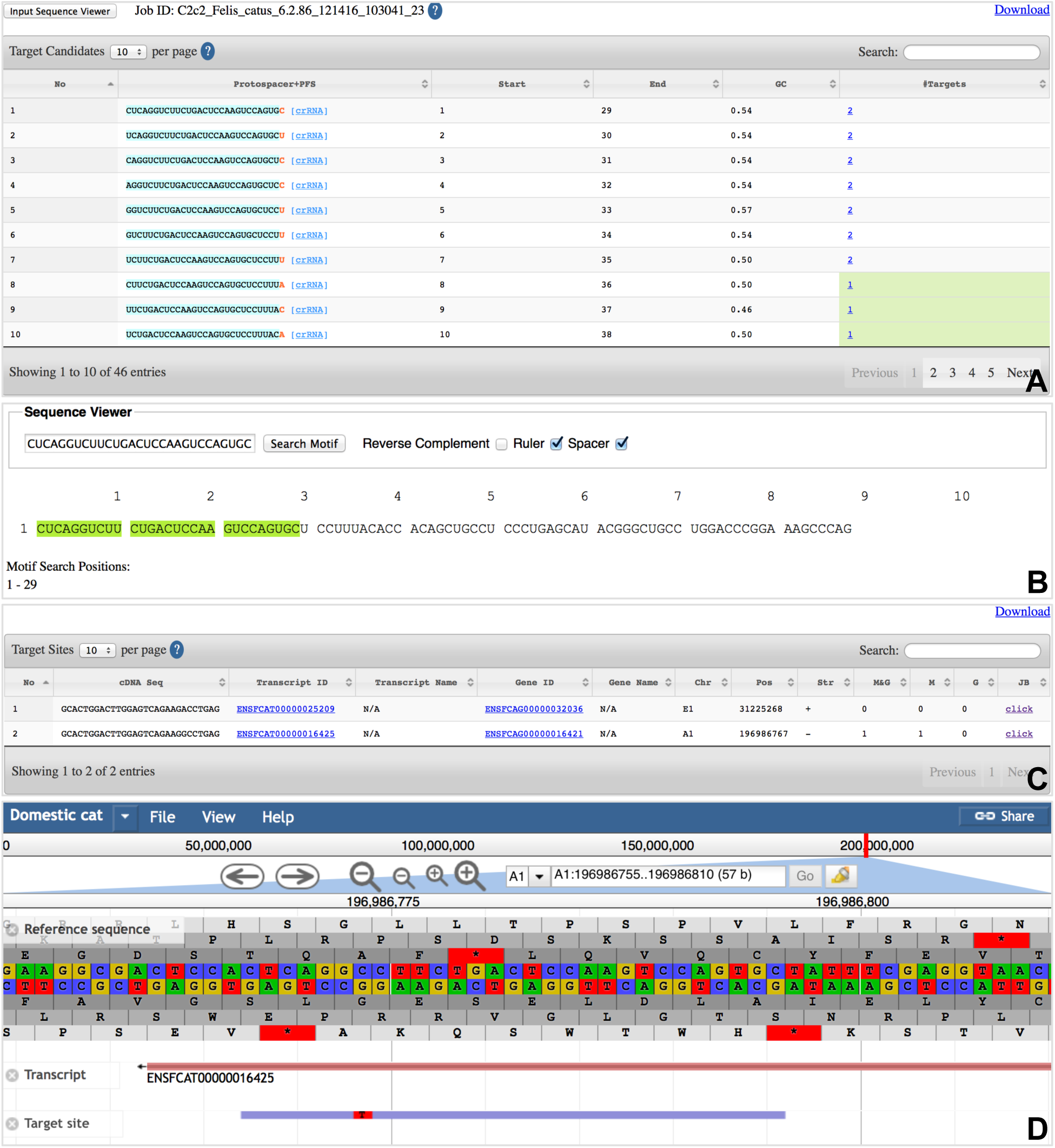
The result web interfaces of CRISPR-RT. (**A**) The detailed information of all the target candidates in the user input RNA sequence. (**B**) User input RNA sequence viewer. (**C**) The detailed information of target sites in the transcriptome for each target candidate. (**D**) Visualization of on- and off-targets with gene and transcript annotations in JBrowse.

**Figure 5.**
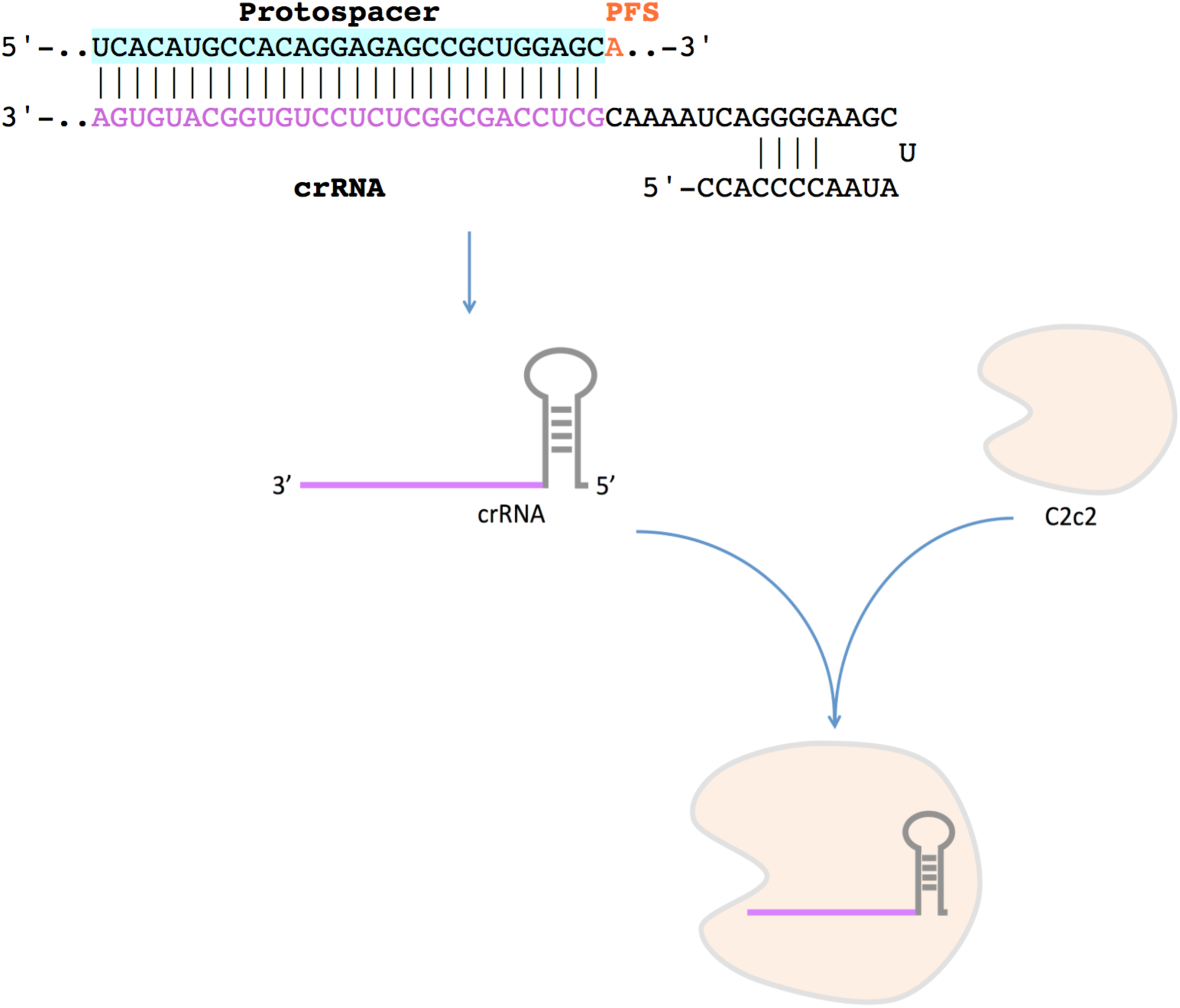
The basic process of crRNA design based on the target candidate sequence (protospacer). The target complementarity sequence of crRNA is labelled in purple color. The stem-loop sequence of crRNA comes from previous research (Abudayyeh *et al.*, 2016). The designed crRNA binds to C2c2 to form a crRNA-C2c2 complex for RNA targeting.

## Discussion

The CRISPR-C2c2 system has been reported as a tool for RNA editing (Abudayyeh *et al.*, 2016). Additionally, CRISPR-C2c2 has already been successfully used for specific RNA knockdown in *E. coli* (Abudayyeh *et al.*, 2016), and it has been applied for RNA detection in human total RNAs (East-Seletsky *et al.*, 2016). The dC2c2, just like dCas9 (Bikard *et al.*, 2013; Qi *et al.*, 2013; Gilbert *et al.*, 2013; Piatek *et al.*, 2015; Maeder *et al.*, 2013; Perez-Pinera *et al.*, 2013; Mali *et al.*, 2013; Cheng *et al.*, 2013; Polstein and Gersbach, 2015; Nihongaki *et al.*, 2015; Zetsche *et al.*, 2015; Gao *et al.*, 2016), also has many potential applications as an RNA-binding protein, such as bringing effectors to specific RNAs to regulate their translation and tracking specific RNAs by fluorescent tag (Abudayyeh *et al.*, 2016). Therefore, the CRISPR-C2c2 system has been viewed as a powerfully programmable tool for current and future RNA editing (Abudayyeh *et al.*, 2016; East-Seletsky *et al.*, 2016; Nainar *et al.*, 2016; Wang and Qi, 2016; Puchta, 2017). However, so far no available software was developed to design crRNAs for the CRISPR-C2c2 system.

In this project, we studied the architecture and features of CRISPR-C2c2 from the bacteria *L. shahii.* By emphasizing the flexible parameters input and the target specificity of crRNAs, we developed CRISPR-RT, the first web-based bioinformatics tool to help biologists design the crRNA for the CRISPR-C2c2 system. Specifically, CRISPT-RT accepts a wide range of parameters such as the number of mismatches or gaps tolerated by off targets, but maintains a simple interface, making it highly flexible for current and future research in CRISPR-based RNA editing and targeting. The powerful alignment tool - Bowtie2 (Langmead and Salzberg, 2012) is used to search all the target sites within the reference transcriptome for each target candidate. Researchers are able to choose the target candidate with minimum off-target effect for crRNA designing. In addition, the on and off-target sites can be validated by data visualization through JBrowse with proper gene and transcript annotations (Skinner *et al.*, 2009).

CRISPR-RT currently covers 10 model organisms including human and can be easily extended to cover other species. The availability of CRISPR-RT will help researchers improve the target specificity of crRNA design for the CRISPR-C2c2 system and empower biologists in CRISPR-based RNA editing and targeting.

## Methods

CRISPR-RT is essentially composed of many web interfaces and a backend pipeline. Web interfaces are implemented by PHP and JavaScript code, which are used to accept user inputs and display the results interactively. The backend pipeline is implemented by Perl code, which is used to process user input data and generate multiple result files.

After setting up proper parameters and clicking the “Find targets!” button in the parameter setting page of CRISPR-C2c2 (Figure 3), all of the parameters stored in PHP code are passed to the main Perl script, which invokes other specific Perl scripts or commands to execute specific functions. As shown in Figure 6, first, the Perl script for target candidate search is called to find target candidates of specified length with the PFS in the input RNA sequence. Second, Bowtie2 (Langmead and Salzberg, 2012) is used to map each target candidate sequence to the reference transcriptome, which is extracted from the genome by RSEM (Li and Dewey, 2011) using Ensembl gene annotation, to search for on- and off-target sites within the transcriptome. Third, the Perl scripts for result filtration based on the input parameters are invoked to filter out the off targets that do not meet the requirements set by users. Next, RSEM is used to convert the transcriptome mapping result to a genome mapping result, which can be displayed properly in JBrowse (Skinner *et al.*, 2009). Then, the main Perl script separates the file storing target sites of all target candidates into many single files that store target sites for each target candidate respectively. The main Perl script also processes the file of target candidates and the files of target sites for each target candidate to generate Ajax format files, which are required by DataTables (a table plug-in for jQuery). After getting all of those files, the PHP and JavaScript code is used to display detailed information of target candidates and corresponding target sites in DataTables. The on- and off-target sites can be visualized in JBrowse.

**Figure 6.**
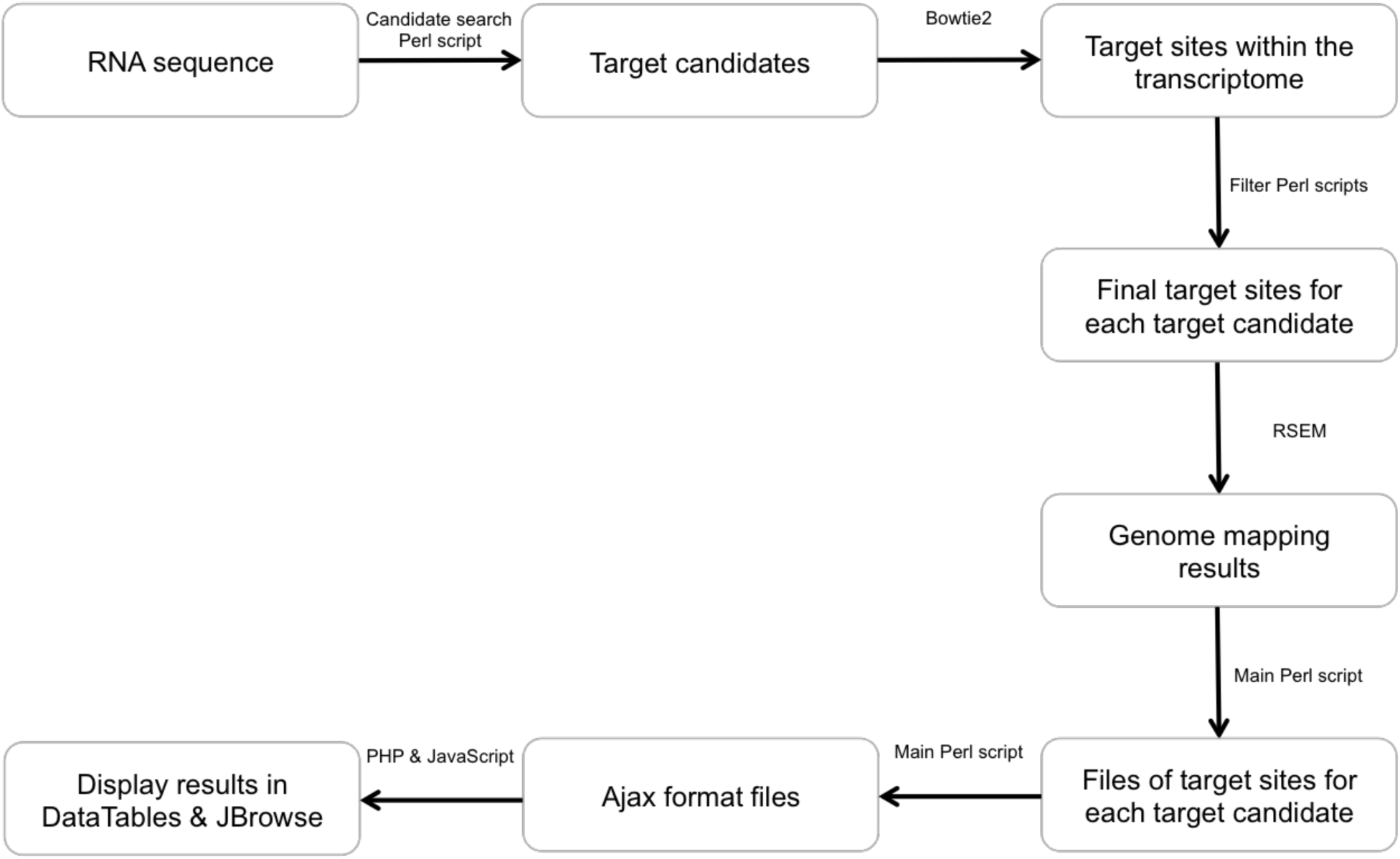
The flowchart of CRISPR-RT backend pipeline that processes user input RNA sequence and displays results in the interfaces. All of input parameters, including the RNA sequence, are passed to the main Perl script. The main Perl script invokes other specific Perl scripts or commands to execute specific functions. Finally, the results will be displayed in DataTables and JBrowse in the web interfaces.

## Acknowledgements

This project is funded by Office for the Advancement of Research and Scholarship (OARS) and Biology Department, Miami University, Oxford, Ohio, USA.

## Authors’ contributions

CL managed and coordinated the whole project. HZ and CL participated in CRISPR-RT design. HZ implemented the backend pipeline in Perl. HZ, ER, and CL participated in web interfaces implementation. HZ, ER, and CL prepared the manuscript. All authors have read and approved the final manuscript.

